# IFT and BBSome proteins are required for *Leishmania mexicana* pathogenicity, but flagellar motility is dispensable

**DOI:** 10.1101/2024.09.13.612850

**Authors:** Tom Beneke, Rachel Neish, Carolina M. C. Catta-Preta, James Smith, Jessica Valli, Ciaran J. McCoy, Andreia Albuquerque-Wendt, Jeremy C. Mottram, Eva Gluenz

**Author notes:** Laboratory of Parasitic Diseases, National Institute of Allergy and Infectious Diseases, National Institutes of Health, Bethesda, MD, USA. Harry Perkins Institute of Medical Research, Perth, WA, Australia. Edinburgh Super Resolution Imaging Consortium, Institute of Biological Chemistry, Biophysics and Bioengineering, School of Engineering and Physical Sciences, Heriot-Watt University, Edinburgh, EH14 4AS, UK. Animal Physiology and Neurobiology, Department of Biology, KU Leuven, Belgium. equal contribution (Tom Beneke, Rachel Neish).

## Abstract

Protists of the order Kinetoplastida possess a single multifunctional flagellum, which powers cellular displacement and mediates attachment to tissues of the arthropod vector. The kinetoplastid flagellar cytoskeleton consists of a nine-microtubule doublet axoneme; further structural elaborations, which can vary between species and life cycle stages, include the assembly of axonemal dynein complexes, a pair of singlet microtubules and the extra-axonemal paraflagellar rod. The intracellular amastigote forms of *Leishmania* spp. build a short, non-motile cilium whose function has remained enigmatic. Here we used a panel of 25 barcoded promastigote cell lines, including mutants lacking genes encoding flagellar assembly proteins, cytoskeletal proteins required for normal motility, or flagellar membrane proteins to examine how these defects impact on their virulence in macrophages and mice. Mutants lacking intraflagellar transport (IFT) protein 88 were severely attenuated indicating that assembly of a flagellum is necessary to allow for *Leishmania* survival in a mammalian host. A similarly severe loss of virulence was observed upon deletion of *BBS2*, a core component of the BBSome complex, which may act as a cargo adapter for IFT. By contrast, promastigotes that were unable to beat their flagella due to loss of *PF16* could establish an infection and only showed a small reduction of parasite burden *in vivo* compared to the parental cell lines. These results confirm that flagellar motility is not necessary for mammalian infection but flagellum assembly and the integrity of the BBSome are essential for pathogenicity.

## Introduction

Protozoan parasites of the order kinetoplastida remain a threat to human health in vast regions of the globe, with an estimated one billion people at risk of infection. Incomplete knowledge of pathogenicity mechanisms and host-pathogen interactions is still considered an impediment to better treatment (Rao, Barrett et al. 2019). Successful completion of their life cycles depends on their ability to establish infections within the blood-feeding arthropod vectors that transmit them, surviving and proliferating in their specific niches within the vertebrate hosts and eventually re-entering a suitable vector. During the course of those life cycles, the flagellum of the kinetoplastid cell serves multiple functions including motility, attachment and sensing (reviewed in (Saez Conde and Dean 2022)). Their motile flagellum has a canonical 9+2 microtubule axoneme adjoined by a kinetoplastid-specific paraflagellar rod (PFR). Eukaryotic flagella (also called cilia) are built from a basal body, docked at the base of the flagellar pocket (FP), by a bi-directional intraflagellar transport (IFT) system that delivers proteins to the tip of the growing flagellum (Absalon, Blisnick et al. 2008, Taschner and Lorentzen 2016). The BBSome, a subcomplex of the IFT system (van Dam, Townsend et al. 2013) acts as a cargo adaptor specifically required for controlling the transport of select membrane proteins into and out of cilia (Wingfield, Lechtreck et al. 2018). Molecular components required for flagellar assembly, structure and beat generation are highly conserved across ciliated eukaryotes (Broadhead, Dawe et al. 2006, Oberholzer, Langousis et al. 2011, van Dam, Wheway et al. 2013, Subota, Julkowska et al. 2014, Beneke, Demay et al. 2019, Vasquez, van Dam et al. 2021). This broad conservation and advanced understanding of many of the key mechanisms that underpin ciliogenesis and organelle function allows for targeted engineering of parasite cell lines with specific deficiencies in flagellum assembly (by blocking IFT) or movement (by removal of cytoskeletal proteins that generate or modulate the flagellar beat), to discover how the flagellum contributes to the parasites’ fitness in the insect vector and test its contributions to pathogenicity.

Genetic studies on *Trypanosoma brucei* have revealed complex dependencies between cell morphogenesis and flagellar assembly during the cell cycle (Kohl, Robinson et al. 2003, Sunter, Webb et al. 2013, Sunter, Varga et al. 2015) and established a vital role for motility in sustaining an infection in the mammal and for completion of its life cycle in the insect vector. *T. brucei* bloodstream forms are particularly sensitive to perturbations of flagellar motility: downregulation or mutation of proteins required for normal flagellar beating lead to rapid and catastrophic failures of cell division and cell death *in vitro* (Broadhead, Dawe et al. 2006, Ralston and Hill 2006), and clearance from infected mice *in vivo* (Griffiths, Portman et al. 2007, Shimogawa, Ray et al. 2018). In tsetse flies, deletion mutants for the axonemal dynein intermediate chain DNAI1 that lost their ability to swim forward (Branche, Kohl et al. 2006) were unable to migrate from the gut to the salivary glands (Rotureau, Ooi et al. 2014).

*Leishmania* promastigotes in the sand fly have a long motile flagellum, which shares fundamental mechanisms with the motile *T. brucei* flagellum. One important difference is the fact that the promastigote flagellum is not attached to the *Leishmania* cell body beyond the exit point from the flagellar pocket, which alleviates the *Leishmania* cell from the absolute dependency of cell division on flagellum growth that exists in *T. brucei* (Sunter, Moreira-Leite et al. 2018). Large-scale phenotyping of *Leishmania* flagellar mutants showed that assembly of a functional motile flagellum was vital for passage through sand flies (Beneke, Demay et al. 2019). Deletion of the central pair-associated protein PF16 produced morphologically normal looking promastigotes, that were paralyzed (Beneke, Madden et al. 2017) and incapable of reaching the thoracic region of the sandfly midgut (Beneke, Demay et al. 2019). Removal of intraflagellar transport (IFT) proteins IFT88 or IFT140 (Sunter, Moreira-Leite et al. 2018, Beneke, Demay et al. 2019) produced viable aflagellate promastigotes capable of productive cell division *in vitro* but unable to persist in sand flies (Beneke, Demay et al. 2019).

The importance of the promastigote flagellum in the infection of mammalian host cells is less clear. Unlike e.g. apicomplexan parasites, where the actin-dependent motility of the parasite drives host cell invasion (Dobrowolski and Sibley 1996, Frenal, Dubremetz et al. 2017), *Leishmania* are thought to rely fully on the host cell’s own mechanisms of phagocytosis to gain entry (Chang 1979, Walker, Oghumu et al. 2014). Studies on polarized engulfment of the promastigotes and the importance of the parasite flagellum yielded mixed findings: While some studies suggested the flagellum may be involved in establishing contact between the parasite and the host cell (Zenian, Rowles et al. 1979, Forestier, Machu et al. 2011), others found that *Leishmania* enter their host cells with their posterior ends first (Courret, Frehel et al. 2002), or with both ends (Chang 1979, Pearson, Sullivan et al. 1983, Rittig, Schroppel et al. 1998). Metacyclic promastigotes, which are pre-adapted to survival in the mammalian host, were seen to exhibit a characteristic “run and tumble” movement, which switched to a faster straighter mode of swimming when presented with human macrophages *in vitro*. This led to the hypothesis that chemotactic sensing promotes active movement of the parasite towards its host cell (Findlay, Osman et al. 2021). Moreover, flagellar beating inside the host cell was reported to trigger host cell response pathways that benefited parasite survival (Forestier, Machu et al. 2011). However, freshly deposited *Leishmania* promastigotes observed by intra-vital 2-photon microscopy were found to be quite immotile in the skin (Peters, Egen et al. 2008) and flagellar beating was shown not to be required for phagocytosis of *Leishmania* (Rittig, Schroppel et al. 1998). To what extent the motile flagellum contributes to infection success has not been answered conclusively. Nor is it certain what function is served by the radical remodelling the flagellum that occurs when metacyclic promastigotes differentiate to amastigotes: within the first 24 hours of experiencing the differentiation signal, the long motile flagellum is remodelled to a short nonmotile 9v type axoneme (Wheeler, Gluenz et al. 2015). Its structural similarity with primary cilia suggested it may act as a sensory organelle (Gull 2009, Gluenz, Hoog et al. 2010) but additional functions in organising amastigote cell morphology, especially the flagellar pocket which is important for endo- and exocytosis, are also conceivable (Gluenz, Ginger et al. 2010).

Here we used fifteen *L. mexicana* gene deletion mutants with different flagellar defects to test whether possession of a flagellum was essential for infection of mice and whether that flagellum needed to be motile. The effect of perturbing membrane protein trafficking was tested indirectly, through disruption of the BBSome. The results showed that the aflagellate *IFT88* deletion mutants were avirulent, while paralyzed *PF16* deletion mutants were capable of establishing infections and persist in the mice for at least eight weeks. Deletion mutants for the BBSome subunit BBS2 were also avirulent despite possessing a motile flagellum. These results show that flagellar motility is dispensable for the mammalian stage of the *Leishmania mexicana* life cycle but IFT and BBSome proteins are required for pathogenicity.

## Results

### A pooled screen of flagellar mutants indicates that flagellar assembly is important for infection of a mammal but flagellar beating is dispensable

A panel of barcoded gene deletion mutants previously used to test the fitness of mutant promastigotes both in culture and in sand flies (Beneke, Demay et al. 2019) was used to assess which type of flagellar defect has an effect on fitness *in vivo*, in a mammalian host.

This pool of five barcoded parental cell lines, five control knockout mutants and fifteen flagellar mutants with a range of motility defects (Figure 1A, Figure S1A-D, Table 1) was used to infect murine bone-marrow-derived macrophages (BMDMs) and BALB/c mice.

**Figure 1.**
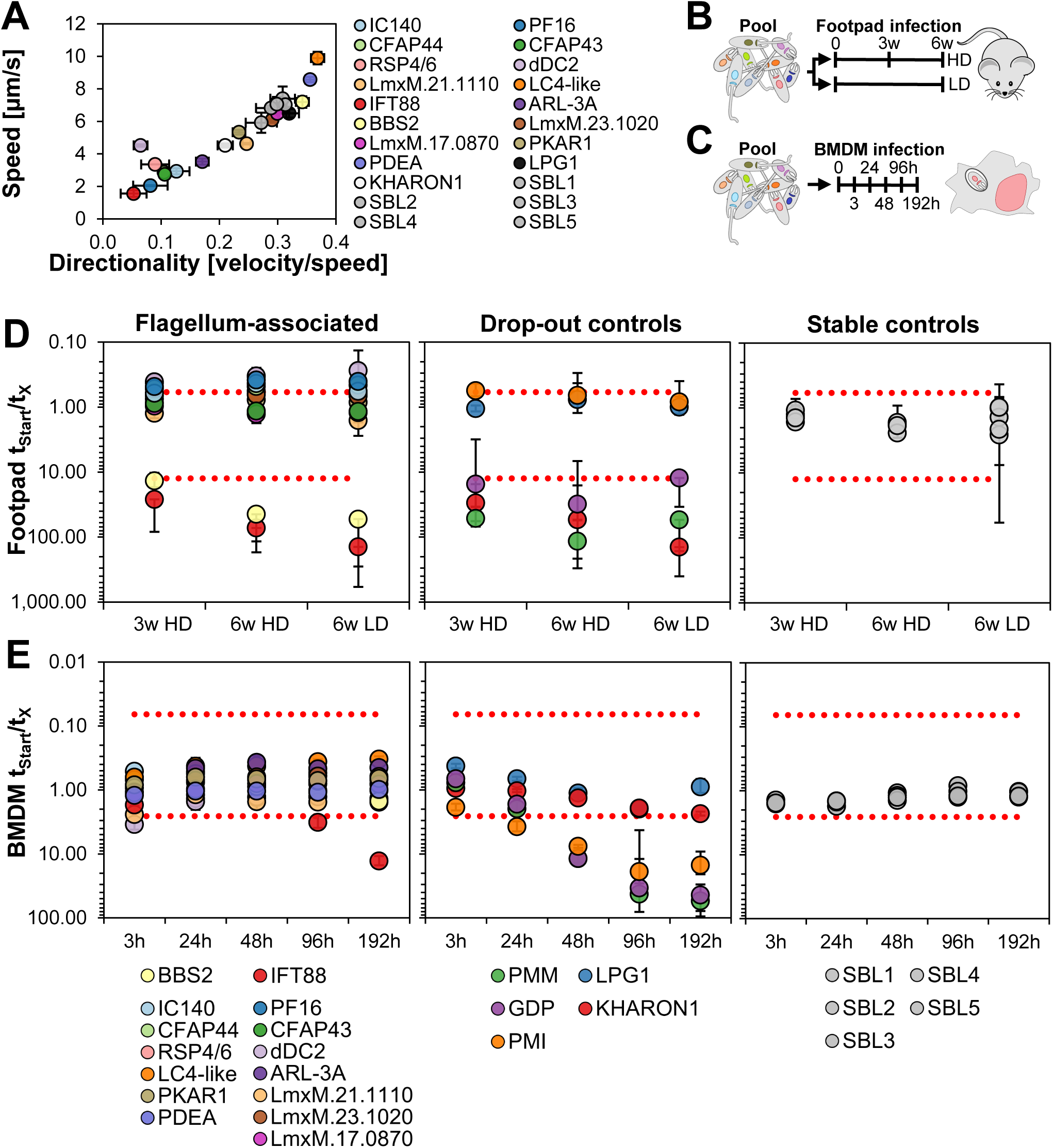
Pooled screen of *L. mexicana* mutants. **(A)** Average speed [µm/s] plotted against directionality [velocity/speed] for the flagellar mutants and the parental controls (SBL1-5) included in the bar-seq screens in BMDMs and mice. The motility data is taken from (Beneke, Demay et al. 2019). The motility of mutants Δ*GDP,* ΔPMI and ΔPMM, which were included in all bar-seq screens as controls, was not measured. **(B)** Schematic of screen design and timeline. At time point 0, pooled stationary phase promastigotes were used to infect mice either with a high parasite dose [HD] or low dose [LD]. **(C)** BMDM infections were started from pools as in (B). Time points of genomic DNA isolation are indicated. **(D)** Bar-seq screen in mice, Y axis: Change in barcode proportions calculated by dividing the normalized read counts at the start of the assay by the normalized read counts at each DNA isolation time point. X axis: DNA isolation time points. Each value represents the average from six infected mice. Error bars show standard deviation between replicates. Red dotted lines indicate two standard deviations above and below the average of “stable control” cell lines. Mutants included in the bar-seq screen have been grouped as follows: mutants with flagellum-associated defects, mutants previously reported to show attenuated virulence (“drop-out controls”) and barcoded parental control cell lines (SBL1-5; “stable controls”). **(E)** Bar-seq screen in BMDMs. Axes and mutant groupings as for (D). Each value represents the average from 3 BMDM samples.

**Table 1.** Cell lines included in pool.

The persistence of null mutants in the pooled infections was tracked over eight days in BMDMs and up to six weeks in mice. This was done by sequencing the barcoded amplicons prepared from DNA samples of infected BMDMs and mouse footpad lesions (Figure 1B). The barcode proportions of the parental cell lines and most mutants remained constant over these observation periods, indicating that most mutants were as fit as the parental controls in the mice and macrophages (Figure 1D,E, Table 2). This included mutants with severe motility defects (e.g. the feebly twitching Δ*CFAP43* and Δ*CFAP44* mutants) and completely paralysed flagella (e.g. Δ*PF16*). In mice, a strong drop-out phenotype was seen for three knockouts of flagellum-associated proteins (Figure 1D, Table 2): Δ*Kharon1,* which lacks a protein required for trafficking of the glucose transporter GT1 to the flagellar membrane, was included in the control group as it had previously been shown to be attenuated *in vivo* (Tran, Rodriguez-Contreras et al. 2013). A similar decline was seen for Δ*IFT88* and for Δ*BBS2*. As expected, the control mutants defective in the pathway leading to mannose activation Δ*PMM,* and Δ*GDP-MP* were also attenuated (Garami and Ilg 2001) but the barcode proportions for Δ*PMI and* Δ*LPG1* did not diminish over time, indicating these mutants had parental-like fitness in this pooled infection assay (Figure 1D, Table 2). The same results were observed in mice infected with a higher parasite dose (2 x10^6^ parasites) or a lower dose (2 x10^5^ parasites). In BMDMs, the control mutants defective in the mannose activation pathway decreased progressively with every time point (Figure 1E). A decline in the proportion of the aflagellate mutant Δ*IFT88* was noticeable at the final time point in BMDMs. The proportion of Δ*BBS2* mutants remained similar to the parental controls after eight days in BMDMs (Figure 1E). The longer-term fate of the *Leishmania* in the BMDMs could not be assessed, a limitation of the *in vitro* system, which highlights the importance of *in vivo* studies to understand pathogenicity mechanisms.

**Table 2.** Sheet 1: Raw and normalized barcode counts for bar-seq screens in BMDM and mice Sheet 2: Calculated ratios (fitness scores) for all bar-screens.

### Paralyzed PF16 mutants are capable of sustaining an infection *in vivo*

The result of the pooled screen suggested a motile flagellum is not required for establishing a successful infection in a mouse. However, if flagellar motility aids the initial infection e.g. by triggering host cell lysosome exocytosis and membrane repair (Forestier, Machu et al. 2011), motility deficient mutants could benefit from the normal flagellar activity of other parasites in the mixed pool. To test whether the Δ*PF16* mutant is able to establish an infection on its own, and persist in a mammal, the Δ*PF16* mutant was used to infect a new set of mice. As a control, a copy of *PF16* was re-introduced into the knockout cell line to generate the *PF16*-addback line (*PF16*-AB; Figure S1E-G). Similarly, addback cell lines were generated for *BBS2* and *IFT88*, to test whether these would rescue the phenotypes of the respective knockout mutants (Figure S1E-G). The Cas9 T7 parental cell line and each knockout and addback cell line were used to infect the right footpad of five individual BALB/c mice. The infections were followed weekly by footpad measurements and parasite burden was calculated for the footpad lesion and lymph nodes, of each mouse.

The footpads of mice infected with the parental cell line showed a progressive increase of lesion size. Lesion sizes from mice infected with the Δ*PF16* mutant initially progressed in a similar way to the parental cell line and to the *PF16*-AB cell line, which had restored PF16 expression (Figure 3A). At the 8-week end point, there was a small difference in lesion size between mice infected with the parental cell line and the Δ*PF16* mutant. Infections with the addback cell line *PF16-AB* restored lesion sizes closer to the values obtained with the parental line (Figure 2A). The parasite burden in the footpad and lymph node was calculated from serial dilutions of extracted tissue. The presence of growing parasites from all samples was noted after three weeks in culture, indicating that live parasites were recovered from mice infected with the Δ*PF16* mutant and the *PF16-AB* cell line from the footpad lesions and from the lymph nodes (Figure 2B). The calculated parasite burden was however lower for the Δ*PF16* mutants, compared to the parental line. These results show that despite being paralyzed, the PF16 knockout parasites were able to establish footpad lesions, persist for eight weeks in mice and remain viable and competent for differentiation to promastigote forms.

**Figure 2.**
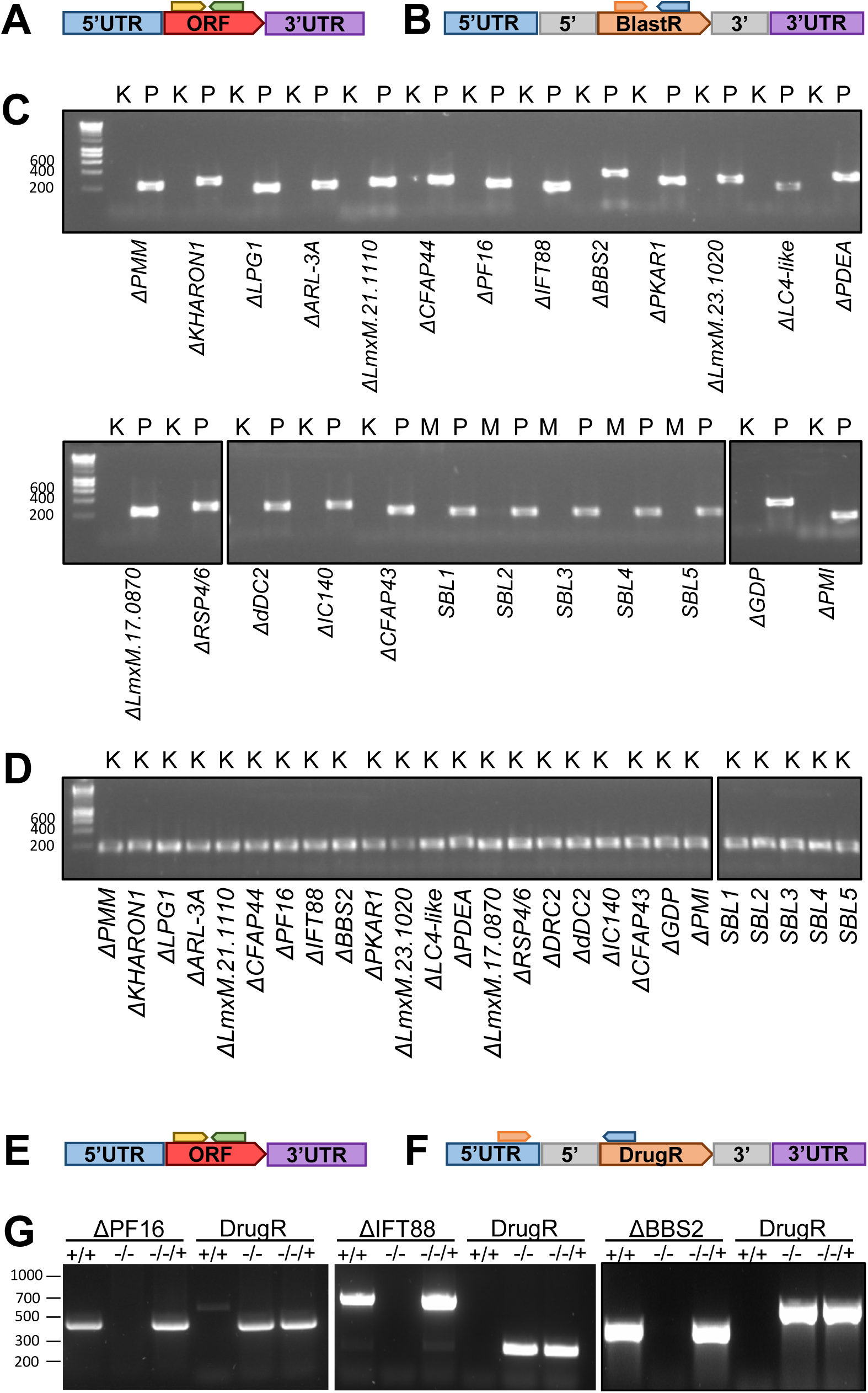
Mouse infections with Δ*PF16,* Δ*IFT88* and Δ*BBS2* Lesion progression and parasite burden were measured for Cas9 T7 parental (+/+), knockout (-/-) and addback (-/-/+) cell lines. (**A, B**) *ΔPF16*, (**C, D**) *ΔIFT88* and (**E, F**) *ΔBBS2*. (**A, C, D**) Measurement of footpad size over time of infection. The average from five mice is shown with error bars indicating standard deviation. P values were calculated using a two-tailed unpaired Students’ t test. (**B, D, F**) Parasite burden measured from serial dilutions from footpad lesions and lymph nodes. The average is indicated. P values were calculated using a Kruskal-Wallis-test with Dunn’s Post-hoc test and Bonferroni correction, comparing knockout and addback against each other (asterisks on line) and against the parental (individual asterisks). (p-value: *<0.05, **<0.0001; ns: non-significant).

### Aflagellate IFT88 mutants fail to establish an infection

The lesion size on the footpads of mice infected with the parental cell line (Cas9 T7) was markedly different compared to Δ*IFT88*, where no lesion growth was seen (Figure 2C). By contrast, the *IFT88*-AB cell line appeared to have the same pathogenicity as the parental cell line, with similar footpad sizes (Figure 2C). Eight weeks after challenge, the mice were culled, dissected and serial dilutions were performed to estimate parasite burden in footpads and lymph nodes after three weeks of culture. Growing parasites were noted for the parental and the IFT88-AB cell line but no parasites were recovered from the footpad lesion and lymph nodes from mice infected with the Δ*IFT88* cell line (Figure 2D). These data show that *IFT88* null mutants are avirulent; they are incapable of causing footpad swelling and likely perish in the mouse. Re-introduction of the *IFT88* gene, which restores flagellar growth (Beneke, Demay et al. 2019), was sufficient to restore pathogenicity.

### The BBSome protein BBS2 is essential for amastigotes

The Δ*BBS2* mutant also failed to produce footpad swelling, in stark contrast to the expected increase of lesion size in mice infected with the parental cell line (Figure 2E). Lesions were noted after infections with the *BBS2-AB* cell line and the footpads grew in size similarly to the parental control infections (Figure 3E). Following serial dilutions of dissected footpad and lymph node tissue after 8 weeks of infection, growing parental and *BBS2-AB* parasites were noted after three weeks in culture and the number of parasites per lesion and lymph node were calculated (Figure 2F). No Δ*BBS2* mutants were recovered from the footpad lesions, nor from the lymph nodes (Figure 2F). These data show that *BBS2* is required for parasite pathogenicity.

The detailed observations on mice infected with individual cell lines support conclusions from the pooled screen, namely that *BBS2* and *IFT88* null mutants are significantly attenuated in their pathogenicity compared to the parental cell line. Virulence was restored by re-introducing the respective gene into the knockout parasites. Footpad measurements and parasite burden calculations for both cell lines indicated they lack the ability to infect the mammalian host. By contrast the *PF16* null mutant was able to infect the mouse and also had the ability to back transform from amastigotes to proliferative promastigotes.

## Discussion

Here we assessed the pathogenicity of *L. mexicana* lines that have been genetically engineered to have distinct deficiencies in flagellum structure or function. The pooled bar-seq screen indicated that perturbations of motility through targeted removal of axonemal proteins (listed in Table 1) had little effect on parasite fitness *in vivo*. Parasites exhibiting a range of motility defects that varied in severity from altered swim speed (increased or decreased, Figure 1A) to complete paralysis remained detectable in infected macrophages for 8 days and in footpad lesions for 6 weeks, with no apparent change in abundance compared to the parental control cell lines (Figure 1D,E). The *PF16* knockout cell line was chosen to verify this finding on the basis of an individual mutant, since its phenotype is already well-characterised in promastigote forms *in vitro* and in the sand fly (Beneke, Madden et al. 2017, Beneke, Demay et al. 2019). Here, despite its inability to move or beat its flagellum, the Δ*PF16* mutant was able to establish footpad infections on its own, clearly demonstrating that a beating flagellum is not required for infection, nor for subsequent survival in an animal host. Between ∼6 and 8 weeks after infection, footpad lesions appeared to plateau in the mice infected with the Δ*PF16* mutant, while still increasing in the control infections. This is consistent with the lower calculated parasite burden at the final timepoint. Whether this indicates enhanced parasite clearance or a reduced parasite proliferation rate *in vivo* is unclear and merits further investigation. The fact that viable *PF16* knockout parasites were recovered after 8 weeks of infection and these were still capable of transforming back to promastigote forms, indicate that the stalling of lesion development is not due to effective parasite clearance.

The finding that motility is dispensable for *Leishmania* in the mammalian host fits well with the naturally occurring remodelling of the axoneme during amastigote differentiation: its 9v axoneme lacks the paraflagellar rod and axonemal dynein motors required for generating a flagellar beat (Gluenz, Hoog et al. 2010, Wheeler, Gluenz et al. 2015). In this respect, *Leishmania* differs from other kinetoplastid pathogens. In *Trypanosoma cruzi* the intracellular amastigotes retain a 9+2 axoneme flagellum shown to beat with ∼0.7 Hz inside host cells (Won, Kruger et al. 2023). Its function remains a matter of speculation (Alves and Bastin 2023) and it would be interesting to test the consequences of interfering with this movement to discover if it is required for pathogenicity. The flagellum of *T. brucei* bloodstream forms is motile and several studies have demonstrated that perturbations of the flagellum lead to rapid cell death *in vitro* (Broadhead, Dawe et al. 2006, Alsford, Turner et al. 2011) and promote parasite clearance from the mammalian host (Shimogawa, Ray et al. 2018). These findings may aid the search for new drugs against African trypanosomiasis (Saez Conde and Dean 2022). For *Leishmania* an essential requirement for flagellar motility has only been shown for promastigotes in their sand fly vector (Beneke, Demay et al. 2019).

Interestingly however, the results with the Δ*IFT88* mutant reported here indicate that assembly of a flagellum, even if paralyzed, or extending only a short distance out of the flagellar pocket, is necessary for successful infection of a mammalian host. IFT88 is a conserved component of the IFT-B complex, which is required for the assembly of microtubule axonemes in most ciliated species. *L. mexicana* promastigotes that lack IFT88 or other IFT proteins can grow as aflagellate cells in culture, albeit with a slightly longer doubling time of ∼10h compared to the ∼7h in the wild type (Sunter, Moreira-Leite et al. 2018, Beneke, Demay et al. 2019). They were however unable to establish a sand fly infection (Beneke, Demay et al. 2019) and the results from the present study show they are unable to infect mice.

Why are the IFT mutants avirulent? A structural role for the flagellum in flagellar pocket architecture could explain this. In promastigotes, the flagellar pocket membrane is attached to the flagellum membrane on one side, leaving a relatively wide opening allowing for fluid access. By contrast, the amastigote flagellar pocket neck closely adjoins the flagellar membrane on all sides, reducing the width of the opening by nearly 10-fold compared to the promastigote FP (Wheeler, Sunter et al. 2016). Disruption of flagellar pocket architecture by deletion of flagellar attachment zone protein 5 (FAZ5) has been shown to attenuate pathogenicity in experimental mouse footpad infections (Sunter, Yanase et al. 2019). While the flagellum is not essential for the formation of the flagellar pocket itself (Sunter, Moreira-Leite et al. 2018) it could serve to protect the flagellar pocket lumen from harmful host factors and contents of the parasitophorous vacuole. The Δ*FAZ5* phenotype would support this hypothesis, since these mutants had a wider FP opening and were slightly more sensitive to the lethal effects of whole mouse serum compared to the parental cell line (Sunter, Yanase et al. 2019). The Δ*IFT88* mutant may be similarly sensitive. Detailed analyses of the morphological consequences of disruption of the IFT-A complex by *IFT140* deletion in *L. mexicana* promastigotes revealed that they still formed a flagellar pocket, which surrounded a short stump of a cilium with a short collapsed 9+0 axoneme that terminated before reaching the flagellar pocket collar (Sunter, Moreira-Leite et al. 2018). While this raised the interesting question of whether assembly of the 9v amastigote cilium is indeed IFT-dependent, or instead triggered by the absence of IFT, the amastigote flagellum is structurally more complex than the rudimentary 9+0 axoneme in IFT deletion mutants. It terminates a short distance beyond the flagellar pocket collar in a bulbous flagellum tip that contains electron dense material of unknown composition (Gluenz, Hoog et al. 2010, Wheeler, Sunter et al. 2016). In a naturally occurring *L. braziliensis* strain with an unusually short flagellum, a normal structural architecture of the flagellar neck region was maintained and this strain remained infection-competent (Zauli, Yokoyama-Yasunaka et al. 2012).

An alternative explanation for how the amastigote flagellum might benefit the parasite is by serving as a point for signal transduction between the parasite and host cell. In mice, IFT and BBS2 deletion mutants exhibited a very similar loss of pathogenicity both in the pooled bar-seq screen (Figure 1D), and in the individual infection experiments (Figure 2). This is compatible with a model where IFT and BBSome functions are linked, as is the case in other cilia where the BBSome acts as a cargo adaptor for IFT, specifically promoting the removal of ciliary proteins (Nachury and Mick 2019). BBSome proteins are highly conserved in ciliated eukaryotes (Hodges, Scheumann et al. 2010, van Dam, Wheway et al. 2013, Ewerling, Maissl et al. 2023). Losing BBSome function was shown to impact on the localisation of ciliary membrane proteins in phylogenetically diverse species (Berbari, Lewis et al. 2008, Lechtreck, Johnson et al. 2009, Domire, Green et al. 2011, Valentine, Rajendran et al. 2012, Datta, Allamargot et al. 2015), with detrimental consequences for ciliary signal transduction. Biochemical studies in *T. brucei* confirmed the direct interaction of BBS proteins in a ∼700 kDa complex in *T. brucei* (Langousis, Shimogawa et al. 2016), and BBS proteins were located at the flagellar transition zone, the gate that controls entry to the ciliary compartment (Dean, Moreira-Leite et al. 2016). In *Lottima passim*, a kinetoplastid parasite of honey bees, the flagellar calcium binding protein FCaBP was found by biotin-proximity labelling to be located close to BBS1 (Yuan and Kadowaki 2024). While IFT-like movement of BBS proteins along the flagellum has not been observed in trypanosomes, this absence of evidence could reflect the technical challenge of detecting low abundance proteins by live cell microscopy (Dean, Moreira-Leite et al. 2016). Flagellar length was reduced upon depletion of Arl6 (BBS3) by RNAi *T. brucei* (Price, Hodgkinson et al. 2012), but flagellum assembly and motility remained normal in the study by (Langousis, Shimogawa et al. 2016) and motility was normal in *L. mexicana* BBSome deletion mutants (Figure 1A). Changes in the *T. brucei* cell surface composition were however noted and the mutants took up less transferrin compared to the wild type (Langousis, Shimogawa et al. 2016). Overall, this evidence is compatible with a model where the BBSome in kinetoplastids acts at the flagellum base to sort specific membrane proteins (Langousis, Shimogawa et al. 2016, Wingfield, Lechtreck et al. 2018), in line with the established function of the BBSome in a range of organisms. What the identity, origin and destination of the sorted proteins is, and whether any have specific functions in the ciliary membrane, is currently not clear, but the process governed by the BBSome is clearly important for the pathogenicity of several kinetoplastid parasite species. The importance of BBSome proteins for the virulence of two other kinetoplastid species has previously been shown, in *T. brucei* (Langousis, Shimogawa et al. 2016) and *L. major* (LmjBBS1 (Price, Paape et al. 2013)). Here a different core subunit of the BBSome, targeted in a different *Leishmania* species, resulted in an equally severe loss-of-fitness phenotype thus replicating and extending the interesting findings by Price *et al*. consolidating the evidence for an indispensable function for the BBSome in *Leishmania* pathogenicity.

## Materials and Methods

### Cell culture

Promastigote-form *L. mexicana* Cas9 T7 (Beneke, Madden et al. 2017), derived from WHO strain MNYC/BZ/62/M379, were grown at 28°C in M199 medium (Life Technologies) supplemented with 2.2 g/L NaHCO_3_, 0.005% haemin, 40 mM 4-(2-Hydroxyethyl)piperazine-1-ethanesulfonic acid (HEPES) pH 7.4 and 10% FCS.

### Genetic modification of L. mexicana

CRISPR-Cas9 gene knockouts were made as described in (Beneke, Madden et al. 2017). Briefly, the oligo sequences from www.LeishGEdit.net were used for the design of sgRNA templates and donor DNA cassettes. The *L. mexicana* Cas 9 T7 parental cell line (Beneke, Dobramysl et al. 2022) containing pRM006 pVY087 was transfected in a single-step transfection with sgRNA templates and donor DNA cassettes derived from pTNeo and pTPuro (BBS2; PF16) or pTNeo and pTBlast (IFT88) and selected with 40 µg/ml G418 and 20 µg/ml Puromycin or 5µg/ml Blasticidin, as appropriate, in supplemented M199 medium at 28°C. For addback cell lines, the coding sequences (CDS) of PF16 (LmxM.20.1400) and IFT88 (LmxM.27.1230) were cloned in plasmid pTAdd. The coding sequence of BBS2 was found to be incorrectly annotated on TritrypDB, starting from a CDS-internal methionine. The full length BBS2 coding sequence was identified by aligning the protein sequences of the conserved BBS2 sequences from *H. sapiens* and *T. brucei* to the translated in-frame ORF upstream of the annotated start codon of LmxM.29.0590. This identified a methionine corresponding to the start codon in the other proteins. This 176 amino acid extension was compatible with the mapped splice acceptor and poly-A sites from *L. mexicana* RNA-seq data (Fiebig, Kelly et al. 2015) and this longer BBS2 CDS (Supplemental Data File) was used to generate the addback plasmid. The sequences of all inserted CDS were checked by Sanger sequencing. The null mutant cell lines were transfected with the relevant addback plasmids and selected with 25 µg/ml Phleomycin in supplemented M199 medium at 28°C.

### Diagnostic PCR for knockout validation

Genomic DNA was isolated as previously described (Rotureau, Gego et al. 2005). Primers for diagnostic PCRs were designed using Primer3 (Koressaar and Remm 2007, Untergasser, Cutcutache et al. 2012) as previously described (Beneke, Demay et al. 2019). Diagnostic PCRs to test for the presence of the target ORF in the putative KO lines and the parental cell lines used in the pooled assays were reported in (Beneke, Demay et al. 2019). All cell lines used in the current study were again tested by PCR before pooling them for the *in vivo* experiments were. Since the SBL1-5 cell lines were generated by replacing the nourseothricin-resistance gene within the 18S rRNA SSU locus with a barcoding cassette, the absence of the resistance gene was confirmed as part of the diagnostic PCR. The cell lines used for individual *in vivo* infections were also tested for the deletion of the *PF16*, *BBS2* and *IFT88* ORF, respectively, as well as for the presence of the reintroduced ORF in each addback cell line. All used primers are listed in Table 3. To show presence of DNA a second PCR reaction was performed to amplify the ORF of blasticidin-S deaminase. SBL barcoded parental cell lines were verified by amplifying the ORF of streptothricin acetyltransferase.

**Table 3.** Primers used for diagnostic PCRs.

### Pooling of mutants for bar-seq screens

For mouse and BMDM infections, sub-pools were seeded at different starting densities, according to their growth rates as promastigotes (Beneke, Demay et al. 2019), to ensure that each mutant in the pool would be equally represented in the stationary phase cultures used for the infections. Sub-pools were seeded at 6 x 10^6^ cells/ml (pool 1; ΔPMM, ΔGDP), 3 x10^6^ cells/ml (pool 2; Δ*IFT88*, Δ*ARL-3A*, Δ*LPG1*, Δ*PMI*), 1.5 x10^6^ cells/ml (pool 3; Δ*IC140*, Δ*PF16*, Δ*CFAP44*, Δ*CFAP43*, Δ*RSP4/6*, Δ*dDC2*, Δ*LmxM.21.1110*, Δ*LC4-like*, Δ*FM458*, Δ*PDEA*, Δ*LmxM.23.1020*, Δ*LmxM.17.0870*, Δ*BBS2*, Δ*KHARON1*) or 1 x10^6^ cells/ml (pool 4; SBL1-5), respectively, and grown for 96 hours. Just before mouse or BMDM infection, the sub-pools were then mixed in proportions to ensure equal representation of mutant lines and a DNA sample was extracted from 1 x10^7^ cells with the Qiagen DNeasy Blood & Tissue Kit.

### BMDM isolation, infection and DNA extraction

Murine bone marrow cells were harvested from femur and tibia of BALB/c mice. After *in vitro* maturation in differentiation medium (DMEM supplemented with 20% L929 conditioned medium, 10 % FBS and 1 % pen-strep) for seven days (with a media change at day 4). BMDMs were cryopreserved until use (in 90% FBS plus 10 % DMSO). Flow-cytometry was used to confirm they were positive for the murine macrophage markers F4/80 (Alexa Fluor488-conjugated antibody, clone BM8) and MAC-1 (Alexa Fluor488-conjugated antibody, clone M1/70), and negative for the granulocyte marker GR-1 (Alexa Fluor647-conjugated antibody, clone RB6-8C5). BMDMs were seeded in 24-well plates (2 x 10^5^ cells per well) and incubated in 1ml differentiation medium for 6h at 37°C, 5% CO_2_. In preparation for infection, the medium was then replaced with fresh pre-warmed DMEM supplemented with 10% FBS and 1% pen-strep and the plates were incubated at 34°C, 5% CO_2_ for 24h before adding the *Leishmania*. Pooled stationary phase *Leishmania* were counted with a haemocytometer and the parasite density was adjusted to 1 x 10^6^ parasites/ml in DMEM + 10% FBS + 1% pen-strep. The medium was aspirated from the macrophages and replaced with 1 ml of the *Leishmania* cell suspension, resulting in a ratio of 5 parasites to one macrophage. After 3h incubation at 34°C, free parasites were washed away with five changes of pre-warmed DMEM with 10% FBS and 1% pen-strep and 1 ml fresh pre-warmed medium was added to all wells except those from the 3h time-point, which were processed for DNA extraction. Remaining plates were incubated at 34 °C, 5% CO_2_ until DNA extraction. DNA was extracted from infected macrophages with the Qiagen DNeasy Blood & Tissue Kit.

### Mouse infections and DNA extraction

Animal work was carried out according to the Animals (Scientific Procedures) Act of 1986, United Kingdom, was approved by the University of York Animal Welfare and Ethical Review Body (AWERB) committee. Thirty female 6 weeks old BALB/c mice were subcutaneously infected in the left footpad with stationary promastigote pools using either 2 x10^6^ parasites (high dose) or 2 x10^5^ parasites (low dose) in 40 μl of sterile PBS. Footpad swelling was recorded (Castanys-Munoz et al., 2012) and increased generally from 1.8 mm to 2.5 mm over 6 weeks. After three and six weeks mice were euthanized by cervical dislocation and DNA was extracted from tissues directly, with the DNeasy Blood and tissue DNA kit (QIAGEN, Cat No./ID: 69506) using the tissue method. Briefly, left footpads were dissected from heel to toes avoiding bones and nails; popliteal lymph nodes were also dissected, samples were immediately weighted, transferred to sterile Eppendorf tubes, and kept at - 20°C overnight. All tissue samples had 15-30 mg and were manipulated under sterile conditions. Samples were incubated for 2h at 56°C in ATL buffer and Proteinase K (Qiagen) to allow complete lyse. DNA was extracted following manufacturer’s instructions.

### Illumina library preparation and sequencing

Sequencing libraries were prepared exactly as described in (Beneke, Demay et al. 2019). Briefly, for each sample, 600 ng isolated DNA was treated with exonuclease VII (NEB) and purified using SPRI magnetic beads. Barcode regions were amplified with custom designed p5 and p7 primers (Life Technologies), containing indexes for multiplexing and adapters for Illumina sequencing. Indices were derived from Illumina Nextera (indices 501-517) and TruSeq (indices RPI1-RPI48) indexing kits (Beneke and Gluenz 2020). Bead-purified amplicons were pooled in equal proportions and the pool was diluted to 4 nM and spiked with 30% single indexed *Leishmania* genomic DNA and 1% PhiX DNA; 8 pM was sequenced using a MiSeq v3 150 cycles kit following the manufactures instructions with paired-end sequencing (2×75 cycles, 6 and 8 cycles index read).

### Bar-seq data analysis

Data analysis followed the process described in (Beneke, Demay et al. 2019). MiSeq raw files were de-multiplexed using bcl2fastq (Illumina). The occurrence of each barcode in the sequencing reads was counted using a bash script (Beneke and Gluenz 2020), searching against the whole database of LeishGEdit barcodes and counting only barcodes with a 100% match to the 17 nt total length. Counts for each barcode were normalized for each sample by calculating their abundance relative to the total number of reads for all 25 barcodes included in the pool. To calculate “fitness” normalized barcode counts in the pooled population before infection were divided by normalized counts at the relevant time point post infection.

### Infection of mice with individual L. mexicana lines and quantification of parasite burden

All cell lines (Δ*BBS2, BBS2-AB*, Δ*PF16*, *PF16-AB*, Δ*IFT88* and *IFT88-AB)* were confirmed to be knockouts via PCR diagnostics. Once confirmed to be null mutants and re-expressors, each cell line, including the parental *L. mexicana* Cas9 T7, were grown to stationary phase promastigotes, left for three days and then used to inoculate the footpad of BALB/c mice at a density of 2.5 x10^6^ per mouse. Virulence of the cell lines was monitored via weekly footpad measurements. The footpad lesion of the control cell lines was allowed to progress to approximately 3 mm. Eight weeks after challenge, the mice were culled, infected footpads and popliteal lymph nodes were dissected as described above and macerated. The tissues were mechanically dissociated and filtered through a 70 μm cell strainer. Homogenates were resuspended in HOMEM supplemented with 20% foetal bovine serum (Gibco) and 1% Penicillin/Streptomycin (Sigma) and serial dilutions were performed. These plates were incubated at 25 °C for approximately three weeks. The presence of growing parasites was noted and parasite burden calculated.

## Supporting information

Supplemental File Protein-sequence

Supplemental Table 1 cell lines

Supplemental Table 2 bar-seq

Supplemental Table 3 primers

## Acknowledgements

We thank Amanda Williams (University of Oxford) for help with Illumina sequencing, Gareth Purvis (University of Oxford) for help with BMDM isolation and Keith Gull (University of Oxford) for access to equipment and helpful comments on the manuscript.

## Supporting Information

**Figure S1.**
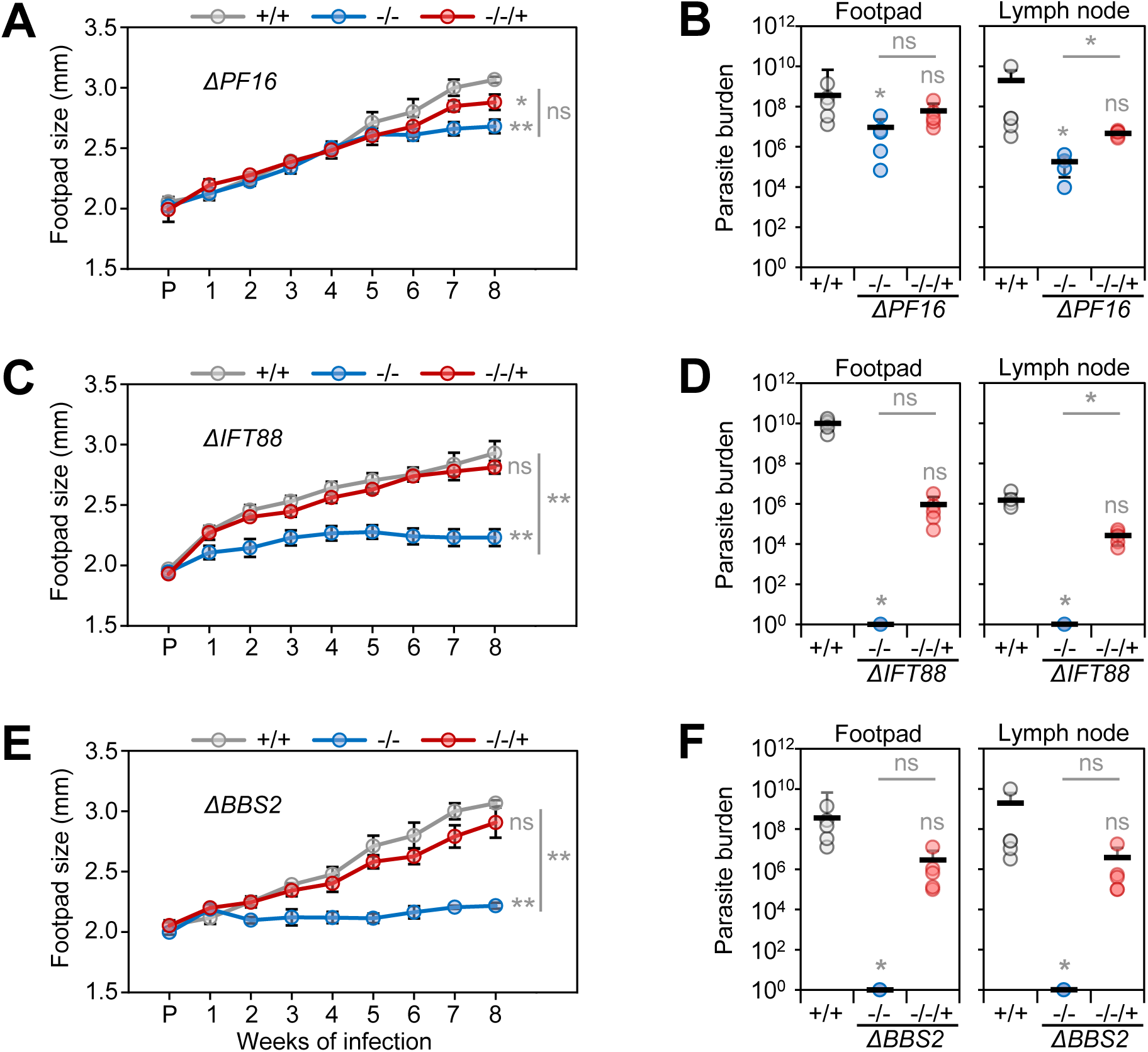
Null mutant PCR validation. **(A, B)** Schematic illustrating diagnostic PCRs to test for the loss of the target ORF in the knockouts used in the bar-seq screens and presence of the integrated blasticidin resistance gene (BlastR). Positions of primers are indicated. **(C)** Results of PCRs testing for the presence of the target ORF. Amplicons were run on an agarose gel. The first lane is the size markers, numbers on the right indicate fragment length in base pairs. K: Putative knockout cell line. P: Parental Cas9 T7 cell line. M: Mutant cell line (SBL1-5). **(D)** Results of PCRs testing for the presence of BlastR in the putative knockout cell lines. **(E,F)** Schematic illustrating diagnostic PCRs to test the cell lines used for individual mouse infections. +/+: Parental Cas9 T7 cell line, -/- knockout cell line, -/-/+ addback cell line. Numbers on the right indicate fragment length in base pairs.

**Supplemental Data File**

The full coding sequence of *L. mexicana* BBS2.

